# Lack of detection of SARS-CoV-2 in British wildlife 2020-21 and first description of a stoat (*Mustela erminea) Minacovirus*

**DOI:** 10.1101/2023.04.28.538769

**Authors:** Ternenge Apaa, Amy J. Withers, Laura MacKenzie, Ceri Staley, Nicola Dessi, Adam Blanchard, Malcolm Bennett, Samantha Bremner-Harrison, Elizabeth A. Chadwick, Frank Hailer, Stephen W.R. Harrison, Xavier Lambin, Matthew Loose, Fiona Mathews, Rachael Tarlinton

## Abstract

Repeat spillover of SARS-CoV-2 into new hosts has highlighted the critical role of cross species transmission of coronaviruses and establishment of new reservoirs of virus in pandemic and epizootic spread of coronaviruses. Species particularly susceptible to SARS-CoV-2 spill-over include Mustelidae (mink, ferrets and related animals), cricetid rodents (hamsters and related animals), felids domestic cats and related animals) and white tailed deer. These predispositions led us to screen British wildlife with sarbecovirus specific qPCR and pan coronavirus PCR assays for SARS-CoV-2 using samples collected during the human pandemic to establish if widespread spill-over was occurring. Fourteen wildlife species (n=402) were tested, including : 2 Red Foxes (*Vulpes vulpes*), 101 Badgers (*Meles meles*), 2 wild American Mink (*Neogale vison*), 41 Pine Marten (*Martes martes*), 2 Weasels (*Mustela nivalis*), 7 Stoats (*Mustela erminea*), 108 Water Voles (*Arvicola amphibius*), 39 Bank voles (*Myodes glareolous*), 10 Field Voles (*Microtus agrestis*), 15 Wood Mice (*Apodemus sylvaticus*), 1 Common Shrew (*Sorex aranaeus*), 2 Pygmy Shrews (*Sorex minutus*), 2 Hedgehogs *(Erinaceus europaeus*) and 75 Eurasian Otters (*Lutra lutra*). No cases of SARS-CoV-2 were detected in any animals, however a novel minacovirus related to mink and ferret alphacoronaviruses was detected in stoats recently introduced to the Orkney Islands. This group of viruses is of interest due to pathogenicity in ferrets. The impact of this virus on the health of stoat populations remains to be established.

## 3. Introduction

Coronaviruses are a large and diverse group of enveloped RNA viruses found in diverse vertebrate hosts. Well-studied mammalian hosts, such as humans and domestic dogs, have multiple coronaviruses of several different subfamilies, although most of those of concern in mammals are members of the *Alpha*- and *Betacoronavirus* genera (Woo, Lau et al. 2012). The SARS-CoV-2 pandemic is thought to have arisen via a spil over from horseshoe bats (the natural hosts of the *Betacoronavirus* subgenus sarbecoviruses), with human infection likely via a ‘liaison host’ such as the raccoon dog (*Nyctereutes procyonoides*) or Malayan pangolin (*Manis javanica*) (Holmes, Goldstein et al. 2021, Crits-Christoph, Gangavarapu et al. 2023, Liu, Lin et al. 2023).

The overwhelming scale of the global human pandemic led to repeated spill overs and onward transmission into other mammalian species such as domestic cats (*Felis cattus*) (Hosie, Hofmann-Lehmann et al. 2021, Goletic, Goletic et al. 2022, Jairak, Chamsai et al. 2022), farmed American mink (*Neogale vison*) (Oreshkova, Molenaar et al. 2020, Domańska-Blicharz, Orłowska et al. 2021, Eckstrand, Baldwin et al. 2021, Hammer, Quaade et al. 2021, Wasniewski, Boué et al. 2023) Syrian hamsters (*Mesocricetus auratus*) (Kok, Wong et al. 2022, Yen, Sit et al. 2022), and the establishment of a new reservoir in North American white tailed deer (*Odocoileus virginianus*) (Hale, Dennis et al. 2022, Kuchipudi, Surendran-Nair et al. 2022). A wide range of other species are either able to be infected experimentally or have been subjects of sporadic case reports of SARS-CoV-2 infection or seroconversion including: cricetid rodents felids (other small carnivores and mustelids primates and bats, reviewed in (Kuchipudi, Tan et al. 2023, Nielsen, Alvarez et al. 2023).

SARS-CoV-2 infection in Muridae such as house mice (*Mus musculus*) and brown rats (*Rattus norvegicus*) may depend on the virus strain: initial studies with the original (Wuhan) strains of the virus failed to infect them (Dinnon, Leist et al. 2020, Shuai, Chan et al. 2021) and field studies failed to demonstrate evidence of infection in wild populations (27 *M.musculus and 97 R.Norvegicus)*. Later variants did however cause infection in laboratory studies (Gu, Chen et al. 2020, Shuai, Chan et al. 2021, Halfmann, Iida et al. 2022, Zhang, Cui et al. 2022) and there have been several subsequent t field reports of sporadic infection of rats (Fisher, Airey et al. 2023, Robinson, Kotwa et al. 2023, Wang, Lenoch et al. 2023).

This study focused on species of wild animals present in Great Britain that were assessed to be of higher risk for SARS-CoV-2 spillover in 2021, when the study was begun. These included, horseshoe bats (subject of a separate report ((Apaa, Withers et al. 2023)) mustelids, small carnivores and cricetid rodents.

Thus far, no infection with SARS-CoV-2 has been reported in Horseshoe bats (*Rhinopholus* spp) in Britain or mainland Europe (Orłowska, Smreczak et al. 2022, Sander, Moreira-Soto et al. 2022, Apaa, Withers et al. 2023) although other related coronaviruses have been detected in these species. Reports in European deer prior to 2022 all failed to detect any exposure (Holding, Otter et al. 2022, Moreira-Soto, Walzer et al. 2022, Wernike, Fischer et al. 2022), however 57% of fallow deer in Dublin seroconverted in early 2022 (Purves, Brown et al. 2023) and sporadic seropositivity in fallow and red deer in Spain in 2021-22 has also been reported (Encinas, Escalera et al. 2023) Wild animal surveillance studies in mainland Europe have indicated sporadic detection in wild mustelids, including by qPCR in wild American mink (*N.vison)*, particularly near farmed mink outbreaks, and one otter (*L. lutra*) (Aguiló-Gisbert, Padilla-Blanco et al. 2021, Padilla-Blanco, Aguiló-Gisbert et al. 2022, Sikkema, Begeman et al. 2022). Serological evidence of exposure has been described in 3/14 pine martens (*M. martes*) and 2/10 badgers (*M. meles*) (Davoust, Guérin et al. 2022), but other studies found no evidence of infection in 48 polecats (*Mustela putorius*), 163 badgers or cricetid and murid rodents (694 *M. glareolus*, 2 *Microtus arvalis*, 27 *M. musculus*, 97 *R. norvegicus* and 8 *Apodemus* species) (Wernike, Drewes et al. 2022, Carmona, Burgos et al. 2023, Zamperin, Festa et al. 2023).

## 4. Methods

### 4.1 Sample collection

A total of 402 animals from 14 species (Table 1) were collected through a network of wildlife researchers and volunteers engaged in wildlife conservation, monitoring or pest control. The majority of samples were collected during the human covid pandemic, with the exception of a small number of historical otter samples. The species targeted were primarily mustelids (Otters *L. lutra*, Badgers *M. meles*, Mink, Pine Martens *M. martes*, Weasels *Mustela nivalis*, Stoats *M. erminea*) or cricetid rodents (Water Voles *Arvicola amphibius*, Field Voles *Microtus agrestis*, Bank Voles *M. glareolus*) with a small number of other species collected opportunistically (Hedgehogs *Erinaceus europaeus*, Common Shrews *Sorex araneus*, Pygmy Shrews *S. minutus*, Red Foxes *V. vulpes*, Wood Mice *Apodemus sylvaticus*). Ethical approval was granted by the University of Nottingham School of Veterinary Medicine and Science Committee for Animal Research and Ethics (CARE), and the University of Sussex Animal Welfare and Ethical Review Board.

**Table 1:** Species and Sample Type Screened for SARS-CoV-2.

| Species | Oral Swab | Rectal swab | Lung sample | Faecal sample | Total number of animals | Sample collection dates |
| --- | --- | --- | --- | --- | --- | --- |
| **Mustelids** |  |  |  |  |  |  |
| Otter | - | - | 75 | - | 75 | Jan 2020-April 2022 (51 samples)<br>April 2017-Dec 2019 (25 samples) |
| Badger | - | - | 101 | - | 101 | April 2021-April 2022 |
| Mink | 1 | 1 | - | - | 2 | Aug 2021-June 2022 |
| Pine Marten | - | - | - | 41 | 41 | March 2020- Oct 2021 |
| Weasel | - | - | 2 | - | 2 | Sept 2021 |
| Stoat | 7 | 7 | - | - | 7 | Dec 2021 |
| **Cricetid Rodents** |  |  |  |  |  |  |
| Water Vole | - | - | - | 108 | 108 | August-Nov 2021 |
| Bank Vole | - | - | 7 | 32 | 39 | Jan-Feb 2022 |
| Field Vole | - | - | 5 | 5 | 10 | Nov 2021-Feb 2022 |
| **Other species** |  |  |  |  |  |  |
| Wood Mouse | - | - | - | 10 | 10 | July 2021-Feb 2022 |
| Common Shrew | - | - | - | 1 | 1 | Jan 2022 |
| Pygmy Shrew | - | - | - | 2 | 2 | Jan 2022 |
| Hedgehog | - | - | - | 2 | 2 | Nov 2022 |
| Red Fox | 2 | 2 | - | - | 2 | April 2022 |
| **Total** |  |  |  |  | **402** |  |

Lung samples were taken from postmortem cadavers submitted to the Cardiff Otter Project (otters) or found dead badgers submitted for tuberculosis monitoring to the University of Nottingham (badgers tested for covid were all confirmed culture-negative for *Mycobacterium tuberculosis* complex infections). Mink and Stoat samples (oronasal and rectal swabs from cadavers) were provided from programmes for invasive species control and were from animals that were live trapped and euthanased (Mink) or lethally trapped (Stoats). Rodent samples were faecal samples from animals live trapped in Longworth or Elliot traps for population monitoring (faecal samples taken from traps or latrine/burrow sites), and from a small number of captive Water Voles from a licenced breeding and release programme. A small number of lung samples were taken from animals found dead. Weasel, Hedgehog, and Shrew samples were from a small number of animals caught as bycatch in Longworth or Elliot traps. Pine Marten samples were all faecal samples (scat) from environmental monitoring. Samples were collected from a variety of British locations (Supplementary Information). Collectors were solicited by social media, notices in game and hunting organisation newsletters, letters in the *Veterinary Record* and contact networks of wildlife organisations working with study participants and were provided with sampling packs including gloves, collection and shipping material and instructions. Samples were collected into RNAlater® before either storage at −20°C or direct shipping to The University of Nottingham, depending on the capacity of the collectors.

### 4.2 RNA extraction, reverse transcriptase (RT) and RNA-dependent RNA polymerase (RDRP) gene coronaviruses generic conventional PCR and envelope gene sarbecovirus-specific real-time PCR

RNA extraction from lung tissue, faecal samples, rectal and oronasal swabs, and cell culture supernatant as positive control, was carried out using the Macherey-Nagel RNA tissue extraction kit as per manufacturer’s instructions. The Wuhan SARS-CoV-2 strain positive control sample used throughout this study was kindly donated by Dr Christopher Coleman (Division of Infection, Immunity and Microbes, School of Life Sciences, University of Nottingham, UK). RT was performed in two steps, using M-MLV-RT and random hexamer primers (Promega) as per manufacturer’s instructions. All cDNA products were stored at −20 °C for conventional PCR.

A generic pan-coronavirus PCR assay published by (Woo, Lau et al. 2005) was used to amplify a 440 bp fragment of the coronavirus RDRP gene using Q5® Hot Start High-Fidelity DNA Polymerase (New England Biolabs cat no: M0493S). Primers were: F: GGTTGGGACTATCCTAAGTGTGA and R: CCATCATCAGATAGAATCATCATA. PCR products were purified using the Nucleospin® extract II kit (Macherey-Nagel) according to manufacturer’s instructions and were Sanger sequenced (Eurofins UK).

Real-time PCR was carried out using the Promega GoTaq® Probe 1-Step RT-qPCR System (Promega) with Sarbecovirus-specific envelope gene primers from bat RNA samples as published by (Corman, Landt et al. 2020). Primers and probes were: F: ACAGGTACGTTAATAGTTAATAGCGT Probe: FAM-ACACTAGCCATCCTTACTGCGCTTCG-BBQ R: ATATTGCAGCAGTACGCACACA

RNA and cDNA quality control was assessed via partial amplification of 108 bp of the beta actin gene using a published conventional PCR protocol (Fischer, Freuling et al. 2014). Primers were F: CAGCACAATGAAGATCAAGATCATC and R: CGGACTCATCGTACTCCTGCTT

### 4.3 High throughput sequencing and genome analyses

RNA sequencing was performed on positive samples by Novogene UK, using the Illumina NovaSeq 6000 platform. Quality filtering and trimming to remove adapters, duplicates and low quality reads was achieved using fastp v0.23.1 (Chen, Zhou et al. 2018). Kraken2 v2.1.2 was used for taxonomic classification reads against the Kraken2 viral Refseq database (Wood, Lu et al. 2019) (retrieved on 9^th^ June 2022). Reads were assembled using the coronaSPAdes option in SPAdes genome assembler v3.15.4 (Meleshko, Hajirasouliha et al. 2021) using default parameters. While CheckV v1.0.1, a fully automated command-line pipeline, was used for identification and quality assessment of contigs, contigs were also queried against the NCBI custom BLASTN (v2.12.0) viral database (Altschul, Gish et al. 1990) (retrieved on 3^rd^ July 2022).

Assembled contigs were indexed, and those that were classified and assessed as complete alphacoronavirus genomes were extracted for downstream analysis using SAMtools v1.16.1 faidx option (Danecek, Bonfield et al. 2021). Assembled genomes were annotated in Geneious Prime® (v.2022.2.2) using NCBI coronavirus reference sequences for minacoviruses and tegacoviruses.

### 4.4 Phylogenetic analysis

Complete coronavirus genomes, extracted RDRP, spike, and nucleocapsid nucleotide sequences from alphacoronavirus genomes assembled in this study, and a total of 22 reference alphacoronavirus genomes (all *Minacovirus* full genomes available and a selection of Refseq or well characterised full length isolates of tegacoviruses with the reference sequence of porcine epidemic diarrhoea virus, PEDV, as an outgroup, supplementary information) were downloaded from NCBI, aligned using Mafft v7.490 (Katoh and Standley 2013). Maximum likelihood phylogenetic trees were reconstructed based on complete coronavirus genomes, and four different genes using IQ-TREE v2.0.7 (Minh, Schmidt et al. 2020), using 1000 ultrafast bootstrap approximations using UFBoot2 within IQ-TREE v2.0.7 to evaluate branch support (Hoang, Chernomor et al. 2018). The ModelFinder function within IQ-Tree was used to select the best fitting nucleotide substitution model for phylogenetic reconstruction (Kalyaanamoorthy, Minh et al. 2017). Phylogenetic trees were visualized and annotated in FigTree v1.4.4 (https://github.com/rambaut/figtree/). Using the same approach trees were also similarly constructed for individual genes (25 Spike and Nucleocapsid genes as the S genes of this group of viruses are known to recombine) and for all available *Minacovirus* partial gene fragments of RDRP (RNA dependent RNA polymerase) and Spike (where there are a lot more sequences available than for other parts of the genome).

## 5. Results

No animal sample tested positive on the *Sarbecovirus*-specific E gene qPCR

Four (out of seven) stoat rectal swab samples (57%) tested positive on the pancoronavirus PCR. Sanger sequencing of PCR products indicated that these were alphacoronaviruses of the *Minacovirus* group. No oral swab samples tested positive from the same animals. All stoat samples in this study were from the same population, and were sourced from the Orkney Islands stoat eradication programme (https://www.nature.scot/professional-advice/land-and-sea-management/managing-wildlife/orkney-native-wildlife-project) and are from the same recently introduced and heavily bottlenecked population. All bar one sample in this study were from adult animals with a mix of male and females. Positive samples were from 1 adult male, 1 juvenile female and 2 adult females.

### 5.1 Illumina sequencing Taxonomic classification, genome assembly

Taxonomic classification using Kraken2 identified reads assigned to other viral operational taxonomic units, however, only reads classified to the Coronaviridae viral family are reported in this study. De novo assembly of datasets from the stoats yielded one full length coronavirus contig of 28.1kb (100% quality, 99.9% completeness) and two partial contigs (7.5kb, 27% quality, 26.6% completeness and 27.9kb, 99% quality and 99.1% completeness) from a further two samples. The remaining sample did not yield any coronavirus contigs. The sequences had closest homology to alphacoronaviruses of the *Minacovirus* group. The complete full genome sequence of the most complete contig has been deposited in Genbank (accession number OP933726).

### 5.2 Genome annotation and organization

Genome annotation demonstrated a typical *Alphacoronavirus* genome organisation consisting of 5’ and 3’ UTRs, a large ORF1ab, encoding sixteen non-structural peptides (nsp1-16) making up about 2/3 of the viral genome, and genes encoding four structural proteins: the spike (S), membrane (M), envelope (E) and nucleocapsid (N) (Figure 1). Accessory proteins included open reading frames with homology to ORF 3c and 7b of feline coronavirus isolates. A number of smaller potential open reading frames within these regions were also present, possibly corresponding to other accessory proteins with homology to the ORF3 or 7 of other *Minacovirus* and *Tegacovirus* isolates.

**Figure 1:**
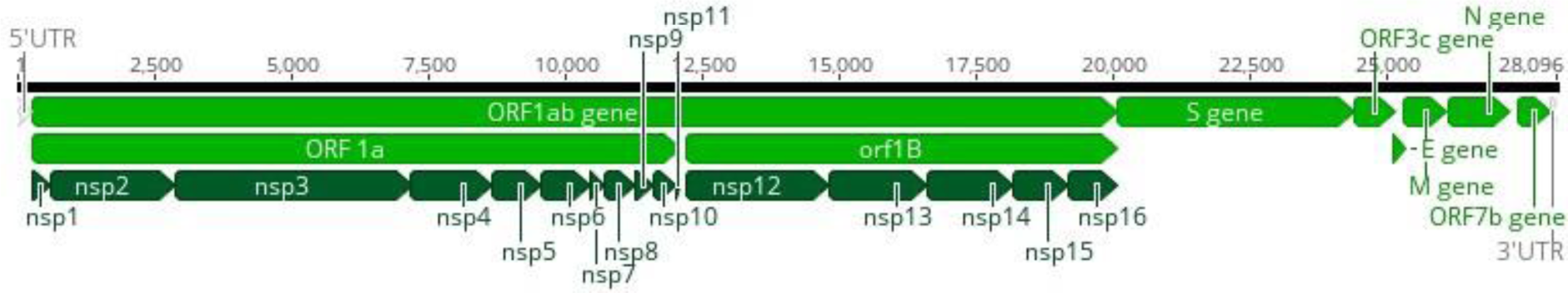
Genome organisation of the stoat sequences derived from this study . Open reading frames are shown in light green, UTR’s in grey and regions with homology to non-structural proteins derived from feline coronavirus ORF1ab are marked in dark green.

### 5.3 Phylogenetic analysis

Results from the maximum likelihood phylogenetic trees drawn using all available complete *Minacovirus* genomes and selected reference sequences for tegacoviruses, demonstrated that the stoat sequence is most closely clustered with ferret isolates (Figure 2). This relationship is consistent across analysis of individual genes (Spike, Nucleocapsid full genes) and in larger phylogenetic trees of all partial fragments of *Minacovirus* RDRP and Spike available in the NCBI database (Supplementary information).

**Figure 2:**
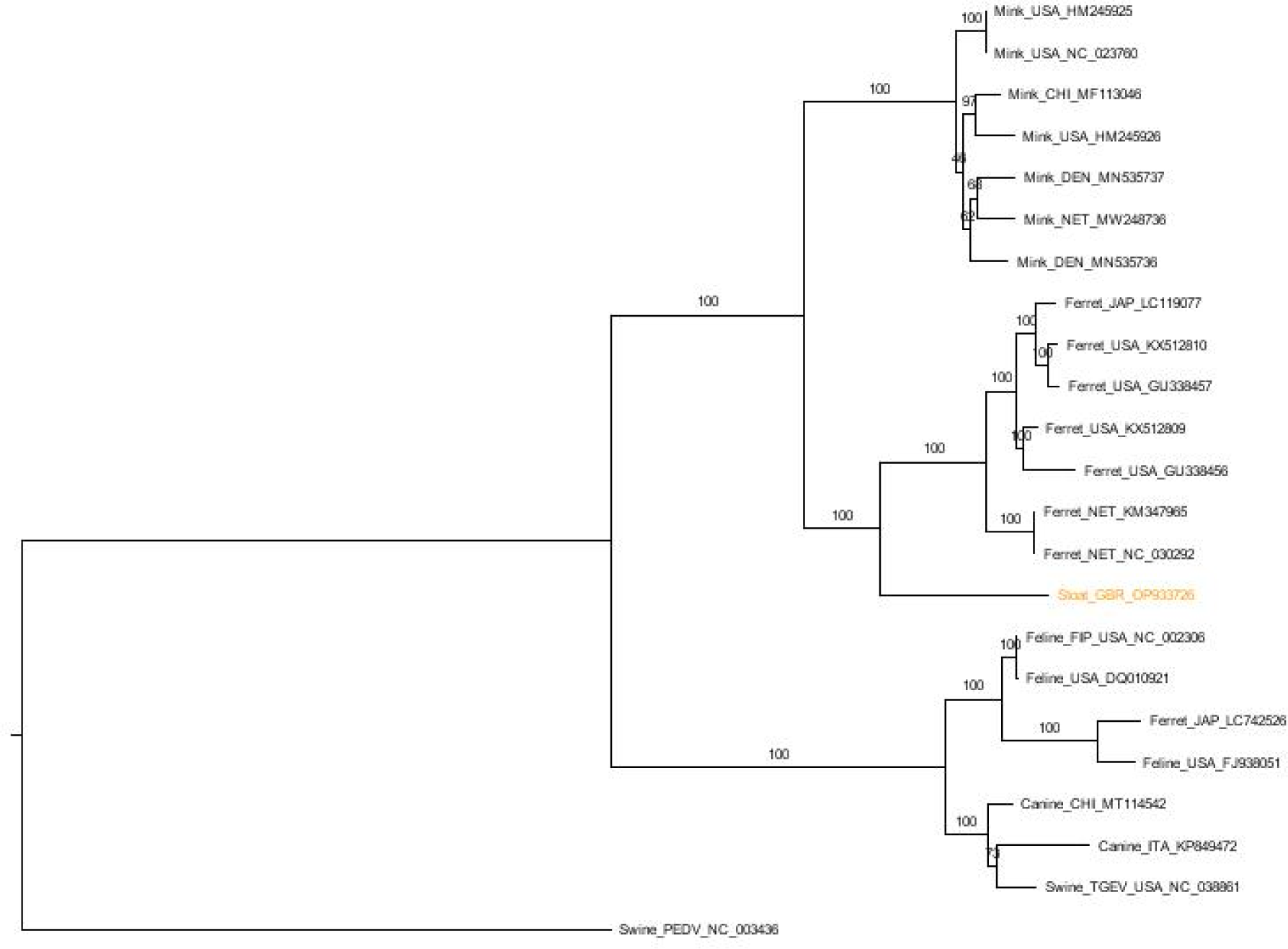
Maximum likelihood phylogenetic tree of full genomes of minacoviruses and tegacoviruses constructed with 1000 bootstrap approximation, rooted on the PEDV coronavirus reference sequence. Twenty three full genomes were included. The sequence from this study is marked in orange. Sequences are named with species of origin, name of virus (if applicable) and a three letter code for geographic origin (GBR=Great Britain, USA= United States of American, CHI=China, JAP=Japan, NET=Netherlands, DEN=Denmark) and Genbank ID. Bootstrap values are shown on nodes

## 6. Discussion

This study found no evidence of widespread circulation of SARS-CoV-2 in our opportunistically gathered British wild carnivores and cricetid rodents in 2021-22. The study was designed to detect an incidence rate of 5% based on our previous studies of alphacoronaviruses in British rodents (Tsoleridis, Onianwa et al. 2016) with a target of 73 samples per species calculated with the online Epitools sample size calculator (Epitools 2020). For some species including otters and badgers, for which there were ongoing post mortem studies of found dead (largely road kill) animals (Sandoval Barron, Swift et al. 2018, Swift, Barron et al. 2021, O’Rourke, Hynes et al. 2022, Thomas, Hailer et al. 2022), this sample size target was readily achieved. This target was also achieved for water voles facilitated by a network of wildlife monitoring and captive breeding for re-introduction (Kirkland and Farré 2021). We were however reliant on volunteer submitters for other species and did not achieve the target levels of fresh samples for, e.g., hedgehogs, foxes, martens and other species. We were also restricted in the type of sample collected for most species as it was primarily non-invasive (faeces or postmortem rectal and oronasal swabs) that could be collected by network members. There remains a possibility that our study did not detect low level circulation of SARS-CoV-2 in the species tested if infection is spatially restricted, that we targeted the wrong tissue sample type or that our samples were too degraded to detect virus with RNA based methods. This is inherent to the study design of a study such as this one seeking to provide the basis for a preliminary assessment with scope to implement a spatially stratified study.

The detection of another coronavirus in this study, however, indicates that the methods used and sample preservation were adequate for viral detection, at least at high prevalence, even in small populations of samples. Our study is also in line with other European wildlife studies indicating absence of widespread SARS-CoV-2 circulation in wild small carnivores and rodents, including wild American mink (Davoust, Guérin et al. 2022, Keller, Peter et al. 2022, Sikkema, Begeman et al. 2022, Villanueva-Saz, Giner et al. 2022, Wernike, Drewes et al. 2022, Carmona, Burgos et al. 2023, Zamperin, Festa et al. 2023). Detection of SARS-CoV-2 in one Eurasian river otter in lung tissue and nasal swabs and detection of a Eurasian badger specific coronavirus in lung tissue from similar post mortem monitoring programmes indicates that lungs are an appropriate target tissue for coronavirus monitoring in those species (Padilla-Blanco, Aguiló-Gisbert et al. 2022, Zamperin, Festa et al. 2023).

SARS-CoV-2 is also readily detected in faecal samples in laboratory studies of rodents and carnivore species as well as field studies of farmed mink (Griffin, Chan et al. 2021, Adney, Lovaglio et al. 2022, Wolters, de Rooij et al. 2022, Wurtzer, Lacote et al. 2022), indicating faecal sample screening is also an appropriate sample for coronavirus monitoring. Faecal samples or rectal swabs are also among the easiest and most acceptable samples for volunteer submitters to collect safely promoting their use in disease surveillance.

Serological testing for SARS-CoV-2 seroconversion would have been a useful addition to this study as PCR based testing used in our study can only detect current viral nucleic acid shedding whereas serology can detect prior exposure to the virus and give a longer term picture of virus exposure. The limitations of volunteer sample submission meant that for many species blood, tissue or body fluids (such as pleural effusion) from which reasonable antibody recovery could be expected were unavailable. Commercial pan-species serology tests SARS-CoV-2 are available and have been used in other studies of wildlife exposure to SARS-CoV-2 (Holding, Otter et al. 2022, Vandegrift, Yon et al. 2022, Wernike, Fischer et al. 2022, Purves, Brown et al. 2023) . These kits are not however validated for all species and we do not know what the cross reactivity is with some of the more divergent coronaviruses detected recently in European wildlife (Zamperin, Festa et al. 2023). Results of these tests are usually confirmed with virus neutralisation assays with false positives a common finding (Davoust, Guérin et al. 2022, Fusco, Cardillo et al. 2023). These caveats make serology testing for an expected low case rate hard to interpret and this work remains for follow up studies.

The stoat alphacoronavirus reported in this study is a novel finding. The stoats all came from an eradication programme on the Orkney Islands, an archipelago off the northeastern tip of mainland Great Britain. They are not native to the islands having been first found in 2010, and cause considerable negative impact on breeding bird populations (Project 2023). Despite the small number of samples collected (7) more than half of them (4) were PCR positive on rectal swabs, suggesting that the prevalence of this virus in this population is high. Wider studies of this population and mainland populations would be warranted to gauge the prevalence of the virus in the larger (and source) populations.

The novel virus we identified in Orkney stoats is a member of the *Minacovirus* subgroup of the Genus *Alphacoronavirus,* and clusters closely to mink and ferret coronaviruses. The ferret and mink viruses have a well described pathogenicity, causing diarrhoea and sometimes a systemic disease syndrome, similar to feline infectious peritonitis in cats (Wise, Kiupel et al. 2010, Vlasova, Halpin et al. 2011, Terada, Minami et al. 2014, Autieri, Miller et al. 2015, Lescano, Quevedo et al. 2015, Doria-Torra, Vidaña et al. 2016, Lamers, Smits et al. 2016, Minami, Kuroda et al. 2016, Wills, Beaufrère et al. 2018). Wider studies of the pathogenesis of this virus in stoats and the impact of this on the species would be warranted. There are multiple reports of the ferret viruses in this group recombining with each other (Lamers, Smits et al. 2016, Minami, Kuroda et al. 2016, Xu 2020) and multiple reports of farmed mink infected with SARS-CoV-2 also being co-infected with minacoviruses at a high prevalence rate (Ip, Griffin et al. 2021, Kwok, de Rooij et al. 2021, Wasniewski, Boué et al. 2023). While recombination between SARS-CoV-2 and minacoviruses has not been observed it is a possibility that warrants monitoring, particularly in farmed mink outbreaks.

Of interest the tegacoviruses (the most closely related alphacoronavirus clade to the minacoviruses) are known for readily recombining and jumping host species, with canine, feline and porcine recombinants in this group reported in multiple separate events (Wang, Ma et al. 2006, Decaro, Mari et al. 2010, Ntafis, Mari et al. 2011, Licitra, Duhamel et al. 2014, Chen, Liu et al. 2019, Pratelli, Tempesta et al. 2022). Canine coronaviruses have also been reported multiple times in multiple locations in people with respiratory disease (Lednicky, Tagliamonte et al. 2022, Vlasova, Diaz et al. 2022, Vlasova, Toh et al. 2022), making this group of viruses of concern for recombination and cross species transmission potential and warranting monitoring.

Sequences in the *Tegacovirus* group have also been reported from raccoon dogs (Wang, Ma et al. 2006, Wang, Tian et al. 2022), suggesting that there could be cross-species transmission between raccoon dogs, domestic cats, dogs, pigs and people. This is of particular concern as this raccoon dogs are known to be able to be infected with and transmit SARS-CoV-2 and are one of the main suspects for the origin of the SARS-CoV-2 outbreak in humans (Freuling, Breithaupt et al. 2020, Crits-Christoph, Gangavarapu et al. 2023, Rao, Parthasarathy et al. 2023).

The *Minacovirus* sequences isolated to date are all associated with mustelids of the genus *Mustela*, subfamily Mustelinae (ferrets, mink, stoats). However there are relatively few studies of coronaviruses in mustelids. That is beginning to be rectified with the publication of SARS-CoV-2 monitoring studies, with a possible *Gammacoronavirus* identified in Chinese ferret badgers (*Melogale moschata*) and, most recently, isolates of a possibly new Genus, *Epsiloncoronavirus,* in Italian badgers (Dong, Liu et al. 2007, Zamperin, Festa et al. 2023). The potential host and geographic ranges of these viruses remain unknown.

Overall this study adds to a growing picture of a lack of widespread SARS-CoV-2 circulation in wild European mammals, other than fallow deer. It has however reported a novel alpha coronavirus of the *Minacovirus* sub-genus in an island population of stoats, adding much needed information on alphacoronavirus diversity in mustelids. The wider disease impacts and epidemiology of this virus in this species is however unknown and requires further study.

## 7. Author statements

### 7.3 Ethical approval

Ethical approval was granted by the University of Nottingham School of Veterinary Medicine and Science Committee for Animal Research and Ethics (CARE), and the University of Sussex Animal Welfare and Ethical Review Board.

## 7.4 Acknowledgements

We are very grateful to the many volunteers who collected samples for this study including Holly Broadhurst, Emma Bryce, Rachel Ewans, Martina Sekirnik, Ellen Bielinksi, Cristobal Castillo, Max Kampmann, John Bruce, Andrea Sartorius, the RSPB Orkney, RSPCA Oak and Furrows, the Vincent Wildlife Trust, Northumberland Wildlife Trust, Thanks also go to the many individuals and organisations who contribute to the otter collection network, including the Environment Agency, Natural Resources Wales, Wildlife Trusts, Local County Ecologists, Trunk Road Agencies, and local mammal groups.

## Footnotes

Company Limited by Guarantee l Registered in England No. 1039582 l Registered Office Charles Darwin House, 12 Roger Street, WC1N 2JU, UK Registered as a Charity: 264017 (England & Wales); SC039250 (Scotland)

