## Supplementary Information for "Lack of detection of SARS-CoV-2 in British wildlife 2020-21 and first description of a stoat (*Mustela erminea) Minacovirus*"

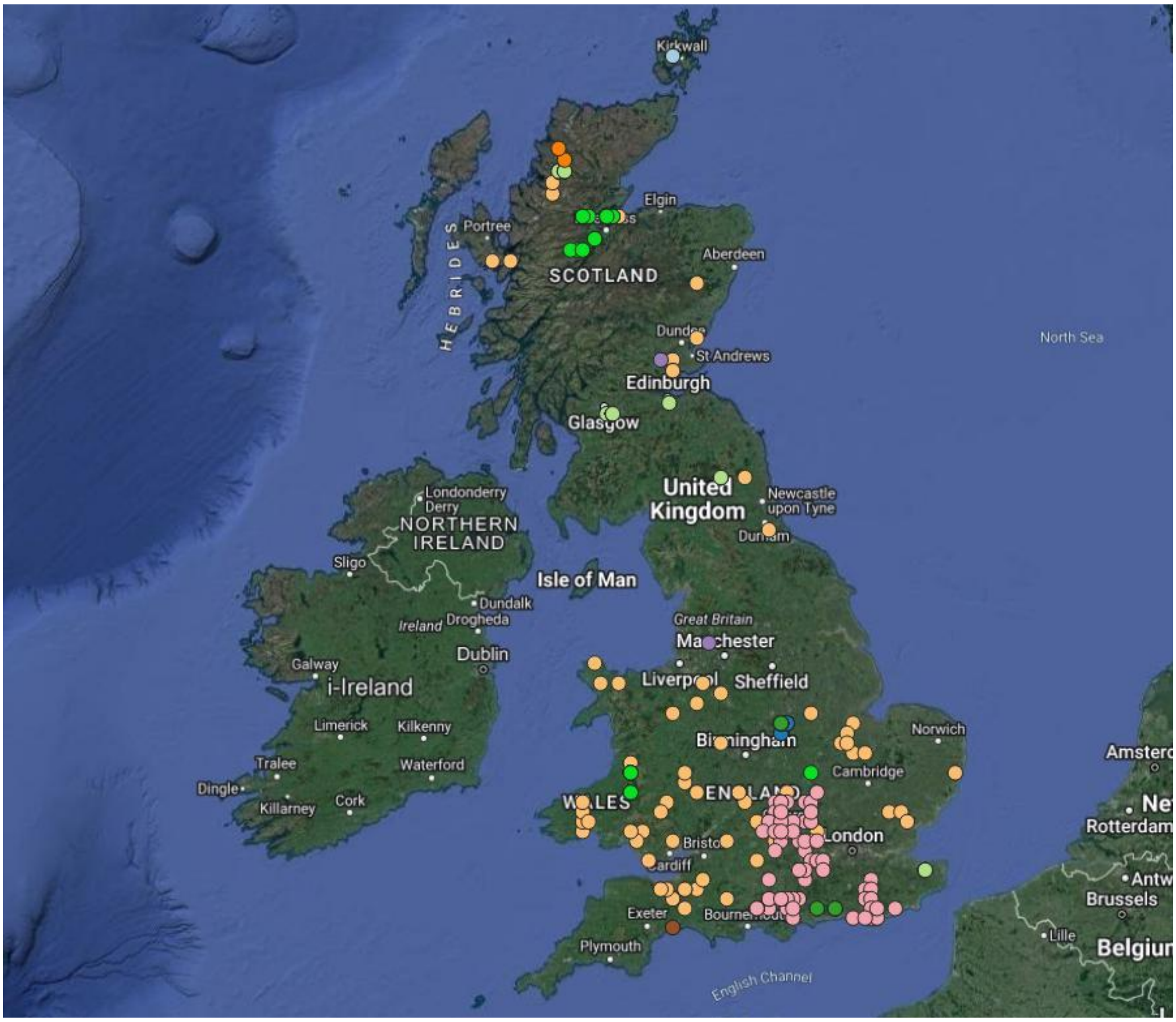


Supplementary Figure 1: locations of Sample collection: Otters= light orange, Hedgehogs= brown, Bank Voles= blue, Badgers= pink, Field Voles=red, Mink= purple, Pine Marten= bright green, Stoat= light blue, Water Vole= light green, Weasel= dark orange, Wood Mice= dark green. Map drawn in QGIS v. 3.30.1-'s-Hertogenbosch

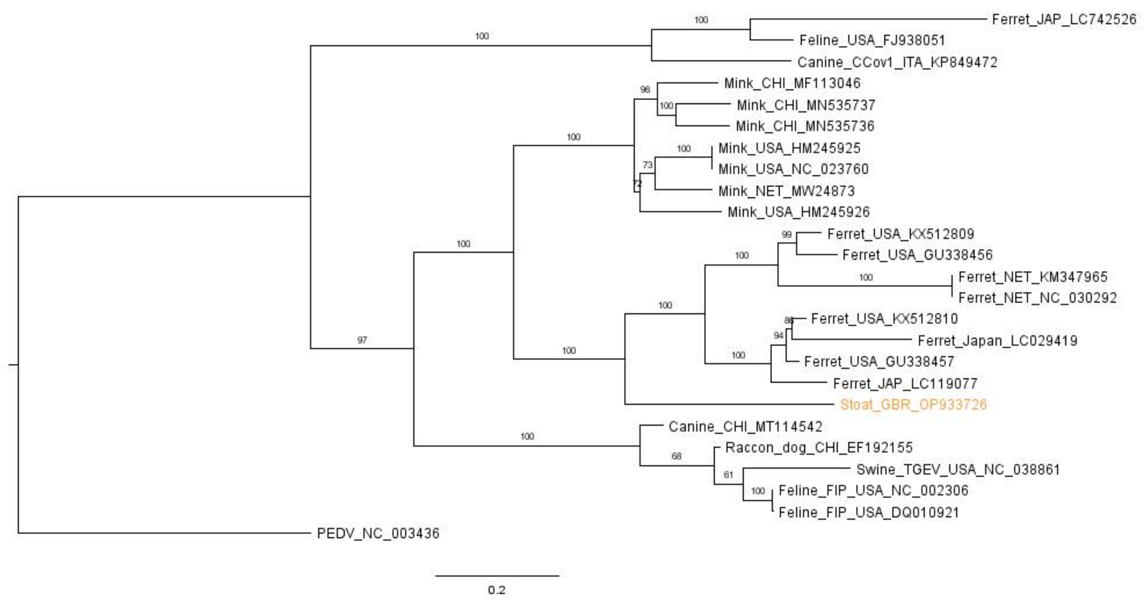


Supplementary Figure 2: Full spike gene minacoviruses and tegacoviruses maximum likelihood phylogenetic tree constructed with 1000 bootstrap approximation, rooted on the PEDV coronavirus reference sequence. Twenty five full gene sequences were included. The sequence from this study is marked in orange. Sequences are named with species or origin, name of virus (if applicable), a three letter code for country of origin (eg GBR=Great Britain) and Genbank ID. Boot strap values are shown on branches.

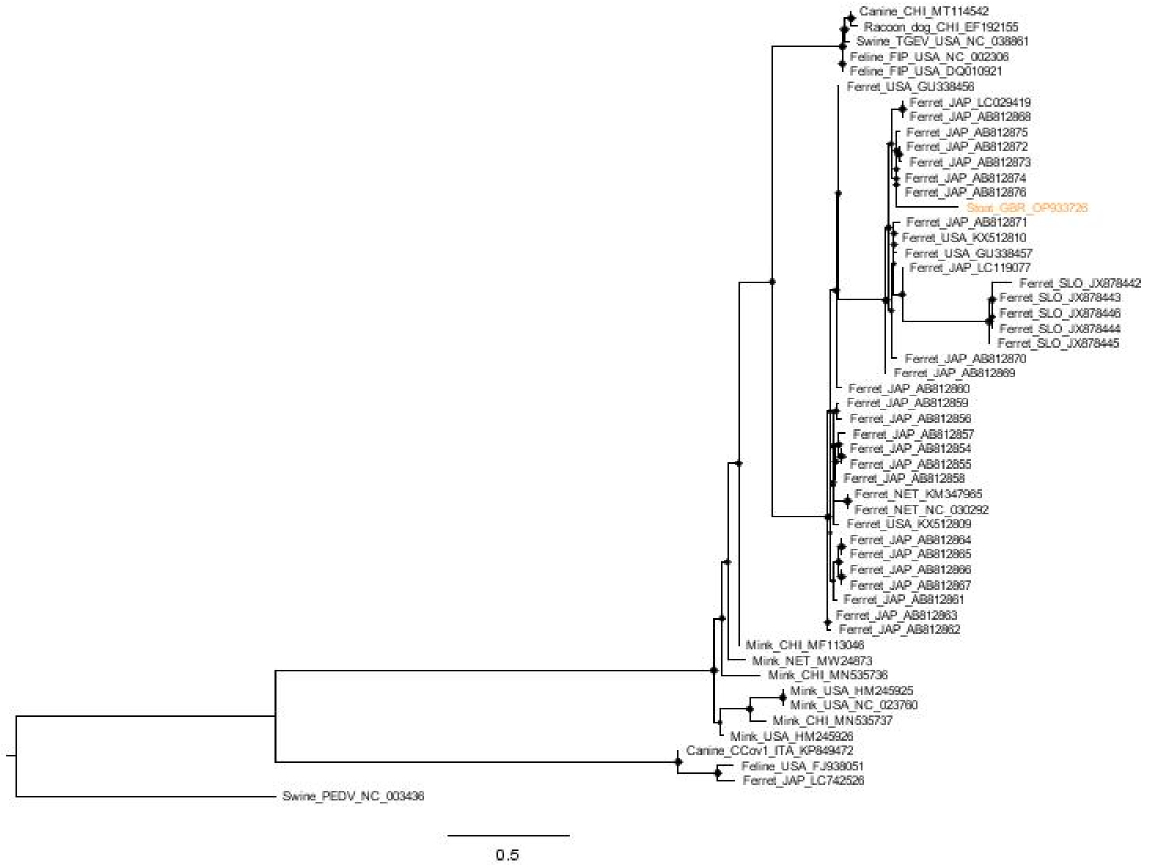


Supplementary Figure 3: Partial fragment (98 bp) spike gene minacoviruses and tegacoviruses maximum likelihood phylogenetic tree constructed with 1000 bootstrap approximation, rooted on the PEDV coronavirus reference sequence. Fifty three sequences were included. The sequence from this study is marked in orange. Sequences are named with species or origin, name of virus (if applicable), a three letter code for country of origin (eg GBR=Great Britain) and Genbank ID. Bootstrap values >50 are shown as diamonds.

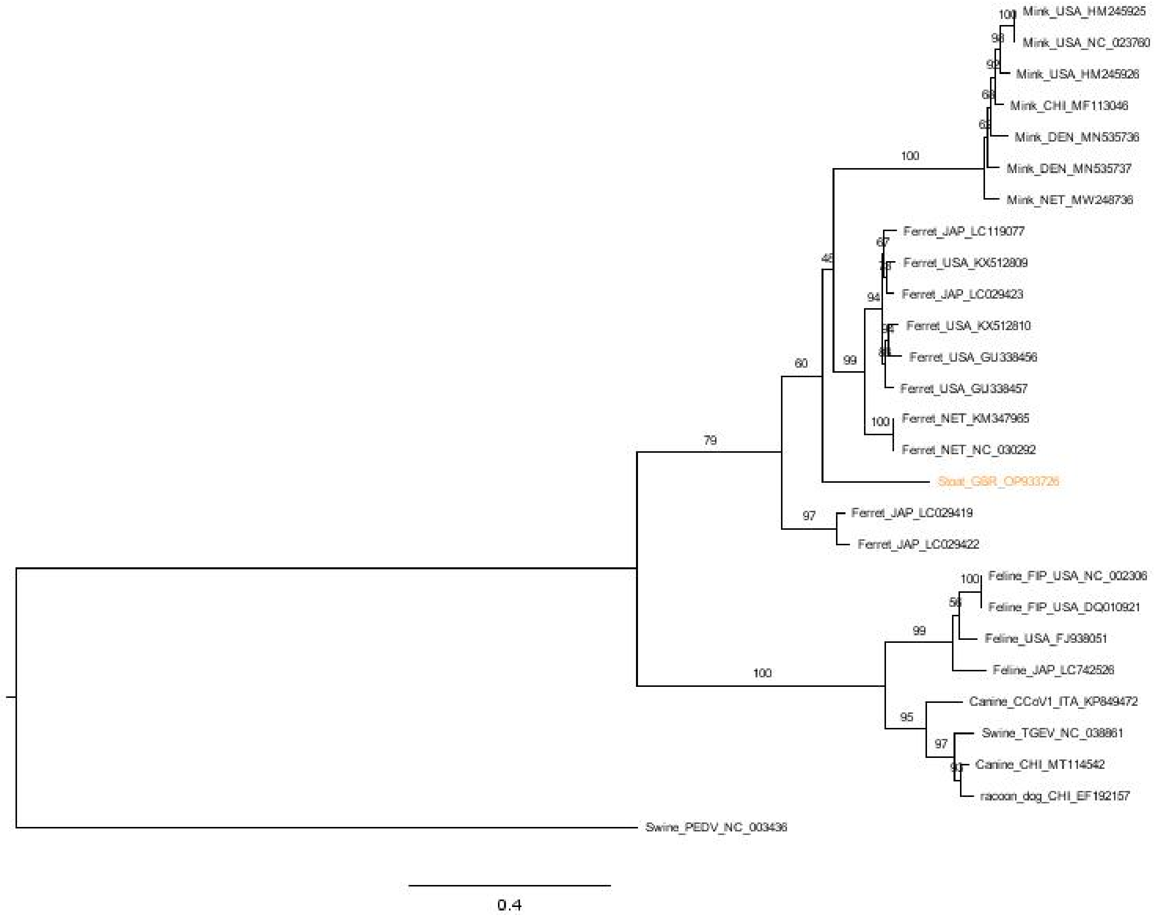


Supplementary Figure 4: Full N gene minacoviruses and tegacoviruses maximum likelihood phylogenetic tree constructed with 1000 bootstrap approximation, rooted on the PEDV coronavirus reference sequence. Twenty seven sequences were included.  The sequence from this study is marked in orange. Sequences are named with species or origin, name of virus (if applicable), a three letter code for country of origin (eg GBR=Great Britain) and Genbank ID. Boot strap values are shown on branches.

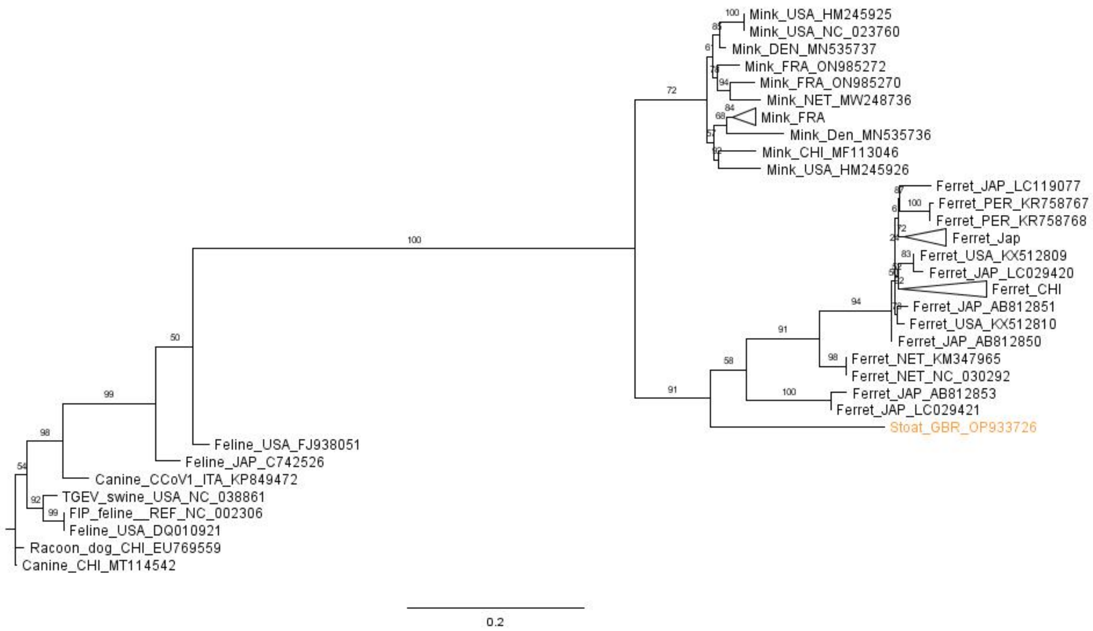


Supplementary Figure 5: Partial fragment (315 bp) RDRP gene minacoviruses and tegacoviruses maximum likelihood phylogenetic tree constructed with 1000 bootstrap approximation. Eighty-nine sequences were included. Mink and Ferret isolates from the same study and country that clustered together phylogenetically have been collapsed for clarity of viewing. The sequence from this study is marked in orange. Sequences are named with species or origin, name of virus (if applicable), a three letter code for country of origin (eg GBR=Great Britain) and Genbank ID. Boot strap values are shown on branches.

Table 1: list of reference genome sequences for full virus trees

| Virus | Genbank ID |
|---|---|
| Porcine endemic coronavirus | NC_003436 |
| Feline coronavirus | DQ010921 |
| Feline coronavirus | NC_002306 |
| Feline coronavirus | LC742526 |
| Feline coronavirus | FJ93805 |
| Transmissible gastroenteritis virus | NC_038861 |
| Racoon dog coronavirus | EQ769559 |
| Canine coronavirus | KP849472 |
| Canine coronavirus | MT114542 |
| Ferret coronavirus | LC119077 |
| Ferret coronavirus | LC215871 |
| Ferret coronavirus | KM347965 |
| Ferret coronavirus | NC_030292 |
| Ferret coronavirus | KX512809 |
| Ferret coronavirus | KX512810 |
| Ferret coronavirus | GU338456 |
| Ferret coronavirus | GU338467 |
| Mink coronavirus | HM245925 |

| | |
|---|---|
| Mink coronavirus | NC_023760 |
| Mink coronavirus | MF113046 |
| Mink coronavirus | HM245926 |
| Mink coronavirus | MN535737 |
| Mink coronavirus | MN535736 |
| Mink coronavirus | MW248736 |
